# The benefits of herbivory outweigh the costs of bioerosion in a eutrophic coral community

**DOI:** 10.1101/2021.06.08.447634

**Authors:** James K. Dytnerski, Katie E. Marshall, David M. Baker, Bayden D. Russell

## Abstract

Herbivores play an integral part in maintaining the health of coral reefs by suppressing the growth of algae and accumulation of sediment and facilitating coral growth. However, in predator-depleted systems where densities of herbivores are unnaturally high, grazing can have detrimental effects on corals through excessive bioerosion. Yet, these benefits and costs are rarely investigated concurrently, especially in eutrophic systems where grazers may play a disproportionate role. We used a year-long exclusion experiment to elucidate the effect of natural densities of the dominant herbivore (the sea urchin *Diadema setosum*) on coral communities in a heavily fished and eutrophic system (Hong Kong, China). To assess benthic community response to grazing, we monitored the survival and growth of three locally abundant coral species (*Pavona decussata, Platygyra carnosus* and *Porites* sp.), algal and sediment accumulation, and bioerosion of coral skeletons across seasons. We found that urchins maintained our experimental coral assemblages, and when excluded, there was a 25 to 75-fold increase in algal-sediment matrix accumulation. Contrary to predictions, there was no general response of corals to urchin presence; *Porites sp.* survivorship increased while *P. decussata* was unaffected, and growth rates of both species was unchanged. Surprisingly, *P. carnosus* experienced higher mortality and bioerosion of up to 33% of their buoyant weight when urchins were present. Therefore, under natural densities, sea urchins clear substrate of algae and sediment, increase survival, maintain growth rates and health of coral assemblages, yet can accelerate the bioerosion of species with porous skeletons following mortality.

## Introduction

Coral reefs are highly diverse and productive systems, providing habitat and resources for 25% of the world’s marine fauna (Molberg and Folke 1999). Herbivores have been well documented to facilitate healthy coral reefs (Hughes 1994, Jackson et al. 2001, Carpenter and Edmunds 2006, Hughes et al. 2007), keeping faster-growing algae in check thereby preventing corals from being overgrown and reducing sediment accumulation (Sammarco 1980, Edmunds and Carpenter 2001, Carpenter and Edmunds 2006, Hughes et al. 2007). Benthic grazers increase reef complexity, providing habitat for fish and invertebrates (Hughes 1994), and increasing habitat suitability for coral larvae settlement by clearing substrate of algal turfs (Sammarco 1980, Davies et al. 2013, Nozawa et al. 2020). Grazing also promotes growth of crustose coralline algae (CCA), which acts as a chemical cue for coral larvae to settle (Heyward and Negri 1999, Harrington et al. 2004, Goméz-Lemos et al. 2018). Furthermore, a healthy grazer community increases ecosystem functions and services (Molberg and Folke 1999), and resilience of reefs to disturbances (Hughes et al. 2007, Graham et al. 2015, Holbrook et al. 2016). The detrimental effects of the loss of grazers on a reef, shifting from of algal dominance to ecosystem collapse, is well documented (Jackson et al. 2001, Hughes, 1994, Bellwood et al. 2004, Carpenter and Edmunds 2006, Mumby et al. 2006). Therefore, with increasing disturbances, such as tropical cyclones and ocean warming-derived bleaching events, replete grazer communities are likely to be essential for coral reefs to persist.

Yet, not all herbivores have the same effects on clearing algae and benefiting corals, as urchins remove more algae than fishes and contribute to higher coral survival rates (Lirman 2001, Mumby et al. 2006, Burkepile and Hay 2008, Reverter et al. 2020). The die-off of the sea urchin *Diadema antillarum* in the Caribbean, during which urchin populations decreased by up to 99% (Lessios et al. 1984, Carpenter 1988), in combination with over-fishing on many reefs (Jackson et al. 2001), led to a shift in reef composition where hard corals were replaced by macroalgae (Edmunds and Carpenter 2001; Carpenter and Edmunds 2006). Conversely, an overabundance of grazers, generally attributed to the removal of predators, can have detrimental effects on ecosystems. Sea otter extirpation from the Aleutian Islands released sea urchins from predation pressure and urchin populations increased, reducing kelp forests to urchin barrens (Estes and Palmisano 1974). In New Zealand, overfishing of spiny lobsters and snapper lead to urchin barrens devoid of most algae; however, this trend was reversed once the top predators returned with the implementation of a no-take marine reserve (Shears and Babcock 2003). In tropical marine ecosystems, an overabundance in herbivores can lead to coral bioerosion. Epilithic bioeroders, like fishes and urchins, target algae growing on the exposed skeleton of corals and scrape away calcium carbonate to eat the algae (Glynn 2015, Alvarado et al. 2017). While coral bioerosion is a natural process - essential for algae removal, coral recruitment, and reef accretion - excessive rates of bioerosion can undercut and topple corals and, on very rare instances, destroy reefs (Bellwood et al. 2004, Glynn 2015). Additionally, eutrophic environments further exacerbate bioerosion rates, as these conditions favour endolithic bioeroders (Barkley et al. 2015, Rice et al. 2020), yet the benefits of herbivory in such systems is less well studied.

The waters around Hong Kong are eutrophic and, as such, conditions have already caused significant loss to local coral abundance and diversity (Duprey et al. 2016, Cybulski et al. 2020). Further, Hong Kong has been extensively overfished and most large predatory and herbivorous fishes found elsewhere in the South China Sea are absent (Cheung and Sadovy 2004, Lai et al. 2016). Consequently, urchins are the primary herbivores which maintain ecosystem functions, as was the case in the Caribbean before the urchin die-off (Jackson et al. 2001). Recently, however, there have been localised reports of large-scale bioerosion on coral communities in Hong Kong caused by the sea urchin *Diadema setosum* (Dumont et al. 2013, Qiu et al. 2014).

While it is well established that urchins play an integral role in maintaining coral reef systems through grazing algae and clearing substrate, few studies quantify the trade-off between herbivory and bioerosion, including coral species-specific effects, particularly in systems in which the key remaining herbivores are urchins. Previous studies have shown that herbivores (fishes and urchins) can increase both growth and survival of corals (Lirman 2001, Hughes et al. 2007, Burkepile and Hay 2008, Burkepile and Hay 2010, Knoester et al. 2019). In contrast, while many studies have estimated urchin bioerosion rates *in situ*, few have simultaneously tested the effects of urchins on both bioerosion and coral growth (Dumont et al. 2013). Therefore, understanding the net benefits and costs of urchins to coral communities could help informing decisions on future coral conservation and restoration efforts.

In this study, we used *in situ* exclusion cages to test the role of a benthic herbivore, the sea urchin *Diadema setosum*, in maintaining coral communities in a highly eutrophic and overfished system. We tested the hypothesis that *D. setosum* would facilitate coral growth and coral survival by removal of algae and sediment. We also assessed the rates of bioerosion at natural urchin densities to test the hypothesis that urchins would not increase bioerosion of live corals. Overall, we expect that urchins will reduce algal-sediment matrix, increase both coral growth and survival, while not increasing coral mortality through bioerosion, thus playing a pivotal role in maintaining corals in a eutrophic system.

## Methods

### Study site

Our urchin exclusion experiment to identify the effects of urchins on coral growth, bioerosion, and survival, and sediment accumulation and turf algal growth was completed at Moon Island (N 22.480878, E 114.333817), Hong Kong SAR, China. The site is located just outside the Hoi Ha Wan Marine Park (HHWMP), a coral habitat protection zone in Hong Kong. It was chosen for the presence of *Diadema setosum* in proximity to coral communities. The site consists of coral communities attached to large rock boulders which formed ideal experimental units for attaching live coral fragments, settlement tiles and coral skeleton cores.

### Experimental design

To test the effects of natural densities of urchins on coral growth, bioerosion, sedimentation and survival, urchins were either excluded (fully caged boulder but left open on the top) or allowed access to plots (urchins present; n = 9 per treatment). In addition, any potential effects of exclusion cages were tested with partial cages which had the bottom and sides open to allow urchins access (procedural control; n = 9; example photos of plots in Fig. S1). All plots included live coral fragments of *Pavona decussata, Platygyra carnosus*, and *Porites sp.*; a skeleton core of *Porites sp.* for monitoring bioerosion; and a 15 × 15 cm terracotta settlement tile (coral sources and skeleton core described below). Items were secured to rock substrate using marine epoxy (Splash Zone A-788 2-part Epoxy, Pettit Paint, Rockaway NJ, USA). These coral species were chosen for their abundance in Hong Kong coral communities (Cybulski et al. 2020). Plots were haphazardly distributed throughout the study site. Cages were rectangular and enclosed ~ 1 m^2^ of substrate. They were built with 2 m long metal rods set ~ 1 m deep into the substrate, wrapped with a plastic-coated wire mesh (2.5 cm mesh squares) approximately 1.5 m high from the substrate. Approximatively 30 cm of mesh was flared out across the substrate at the bottom of the cage to prevent urchins from accessing the cages. The tops of the cages were left open so fishes could enter. Crabs, gastropods, and sea cucumbers were also frequently observed in the cages. Urchins were never observed inside any cages throughout the experiment. Cages were inspected concurrently with monthly surveys, scrubbed periodically to remove algal growth, and repaired when needed.

### Coral growth and survival

Coral fragments were sourced from existing coral nurseries maintained in a shaded 2000 L mesocosm with flow-through seawater at The Swire Institute for Marine Science (SWIMS), Cape D’Aguilar, Hong Kong. Additional samples of *P. carnosus* and *Porites sp.* were collected from Bluff Island (N 22.324386, E 114.353946) and brought back to the coral nursery mesocosm at SWIMS for seven days to recover after fragmentation. Live coral plugs were taken from colonies using a 55 mm diamond tipped hole saw drill bit and carefully pried out with a screwdriver. After the seven-day recovery period, all coral fragments were measured for buoyant weight and then attached to experiment plots. Dead, almost dead, and severely bioeroded fragments of *P. carnosus* were recovered from plots and replaced with new fragments at days 97 and 230 of the 393-day experiment to capture the rate of bioerosion and mortality (Fig. S1). All other coral fragments were collected at the completion of the experiment and buoyant weight re-measured. If corals died, the species, plot type and date was recorded for survival analyses.

### Bioerosion of coral cores

To quantify bioerosion, skeletons of *Porites sp.* from an old museum collection were cored with an 80 mm hole saw drill bit, with the cores then cut into discs approximately 10 mm thick with a rock saw. These coral cores were dried for 48 hours at 60°C and then weighed and 3D scanned for surface area using NextEngine 3D scanner (Santa Monica, CA, USA). Scans were done in 360° mode, with eight divisions, scanning every 45 degrees. Divisions were fused together, producing a solid image using the volume merge function set to a 0.9 resolution ratio in Scan Studio from NextEngine. Surface area was calculated directly by Scan Studio. To monitor summer bioerosion, coral cores were attached in May 2017 and retrieved in December (208-day deployment). New cores were attached in January 2018 for winter bioerosion rates and collected in May (120-day deployment). Retrieved cores were rinsed with fresh water and dried at 60°C for 48 hours then re-scanned and weighed. Mass and surface area data were combined and transformed for analytical purposes to kg · m^−2^ year.

### Settlement tiles

To capture colonization patterns of benthic organisms (algal turf, encrusting algae, and invertebrates) and sediment accumulation trends, settlement tiles were deployed between April and November 2017 (208 days) for summer and November 2017 to March 2018 (92 days) for winter. In addition, photos of each tile were taken *in situ* in July and November (summer) and January and February (winter) for percent cover analysis and colonization over time. At the end of deployment, each tile was gently removed from the substrate, placed into labelled, sealable plastic bags, and brought back to the laboratory and scraped of sediment and algae. The tile scrapings were put into a pre-weighed aluminium weigh boat, dried in a 60°C oven for 48 hours and then re-weighed. Algal-sediment matrix weights were then converted to dry mass by area (g · m^−2^).

Benthic assemblages which established on settlement tiles were quantified using the image annotation platform CoralNet Beta. A 15 × 15-point grid overlaid the settlement tiles, giving 225 points per tile. Each point was evaluated to determine coverage identity. A random sampling of 45 tiles was used to establish a baseline for the automated annotator function. Remaining tiles and points were identified by the CoralNet annotator, and then each point was manually checked and corrected if mislabelled to ensure accuracy of identification. Items identified on the tiles were sorted into five functional groups: bare tile; encrusting algae; fleshy algae; algal-sediment matrix; and sessile invertebrates. The last photo survey immediate before tile retrieval, for both end of summer and end of winter, were used for analyses to compare the affect of urchins on benthic community and sediment accumulation.

### Urchin surveys and monitoring

Urchin density at the study site was quantified monthly using permanent belt transects (25 m × 2 m; n = 3) which covered the area around all experimental plots. Over the length of the experiment, mean density of *Diadema setosum* was 5.4 ± 0.2 urchins per m^2^ (mean ± SE), ranging between 4.3 ± 0.7 and 6.8 ± 1.3 m^−2^, and did not differ among sampling periods (*F* _8, 18_ = 0.27, *p* = 0.97). *D. setosum* was the only urchin species found at the site in substantial numbers, with individuals of *Heliocidaris crassispinus* and *Salmacis sphearoides* seldomly observed.

### Data analyses

To test for the effects of urchin presence on the community composition on settlement tiles, percentage cover of functional groups at the end of each season (summer and winter) were analyzed separately using PERMANOVA and visualized with nonparametric multidimensional scaling (NMDS) plots. All data were square root transformed and put into a Bray-Curtis distance matrix using 999 permutations. NMDS ordination plots were generated using Bray-Curtis distances on two dimensions. Both analyses were conducted in R Studio using the Vegan package (Oksanen et al. 2019). Dry biomass of communities on settlement tiles were compared among experimental treatments using one-way ANOVA. To evaluate the effect of urchin presence on coral survival, we used survival analysis, performed in R Studio with the package survival (Therneau 2020). For *Porites sp*., we performed a next plus one transformation to account for 100% survival in the urchin present treatment and added an artificial death to each treatment. Accelerated Failure Time models were also run to test the effect of urchins on coral survival but found similar results to the Survival Analyses and were thus not included here. The effect of urchins on growth or erosion of coral fragments was tested using a one-way ANOVA for each species. Data were transformed to the percent change per year of each fragment as fragment sizes varied within and among species. Only corals that were still alive at the end of the experiment were included in growth analyses. To investigate the effect of urchins on bioerosion of coral cores, we ran one-way ANOVAs, with summer and winter seasons analyzed separately. All statistical analyses were performed in R Studio (version 3.6.1).

## Results

The presence of urchins caused different benthic communities to develop on settlement tiles in a season-specific way (Fig. 1). Communities where *D. setosum* were present, both open and partial cage plots, were centred around CCA and benthic microalgae (Table S1 for definitions) in the summer, and microalgae and bare tile in the winter, whereas communities when urchin were excluded centre around algal turf and serpulid worm sediment matrices in the summer, and serpulid worms and sepulid worm sediment matrices in the winter (Fig. 1A, B.).

**Figure 1.**
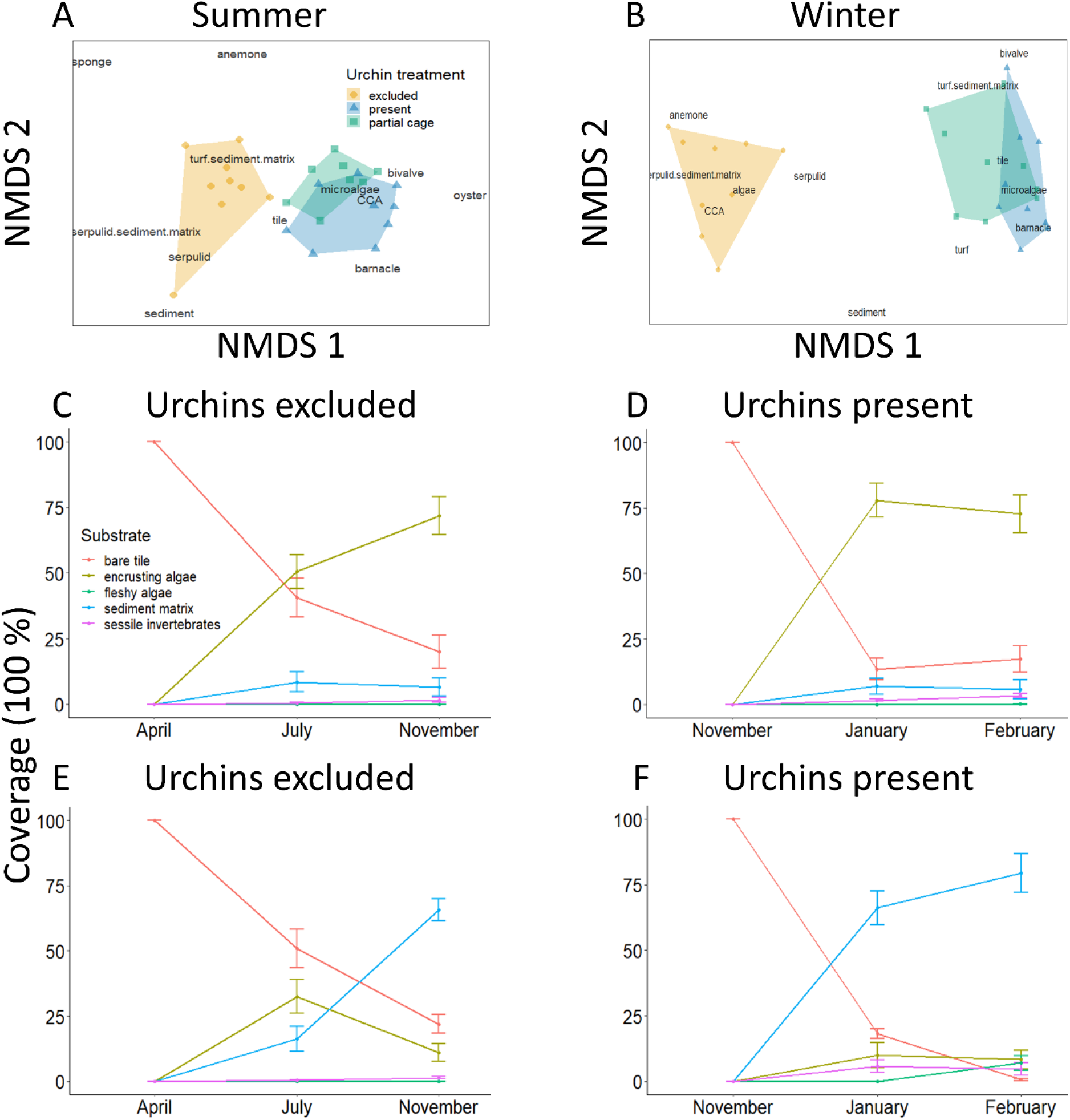
Difference in the final benthic substrate community makeup under different urchin treatments at the end of (A) summer and (B) winter (NMDS plots with stress = 0.10 and 0.08, respectively); and benthic assemblages when urchins were present (C, D) and excluded (E, F) in both summer (C, E) and winter (D, F). Settlement tiles started with 100 % bare surface. Values are mean ± SE.

When benthic taxa are combined into functional groups (Table S1), the presence of urchins reduced the coverage of algal sediment matrix by 60% and increased the cover of CCA by 60% by the end of the summer (Fig. 1C, E; *F*_2, 23_ = 13.01, *p* < 0.001). There was a divergence of benthic community composition at the end of the winter, with an even stronger influence of urchins on the community (Fig. 1B). When urchins were present, CCA coverage was 65% greater than without urchins by the end of the season. Meanwhile, there was a decrease in algal-sediment matrix by 75% (Fig. 1E and F; *F*_2, 24_ = 45.76, *p* < 0.001).

When urchins were excluded, algae and sediment accumulation (algal-sediment matrix) was 29 times greater during the summer and 80 times greater during the winter compared to when urchins were present (Fig. 2; *F* _2, 23_ = 8.59, *p* < 0.002; *F* _2, 24_ = 124, *p* < 0.001). During the summer, there was 22.3 ± 6.9 g · m^−2^ of algal sediment accumulation when urchins were excluded from the plots but < 1 g · m^−2^ when urchins were present. This pattern was exacerbated during winter, with an average of 1027.7 ± 90.5 g · m^−2^ accumulated when urchins were excluded, compared to 12.8 ± 2.0 g · m^−2^ when urchins were present. There were no differences in the mass of algal-sediment matrix between the partial cage treatment and when urchins were present, for either summer or winter (Tukey HSD: *p* = 0.98 and p = 0.99, respectively).

**Figure 2.**
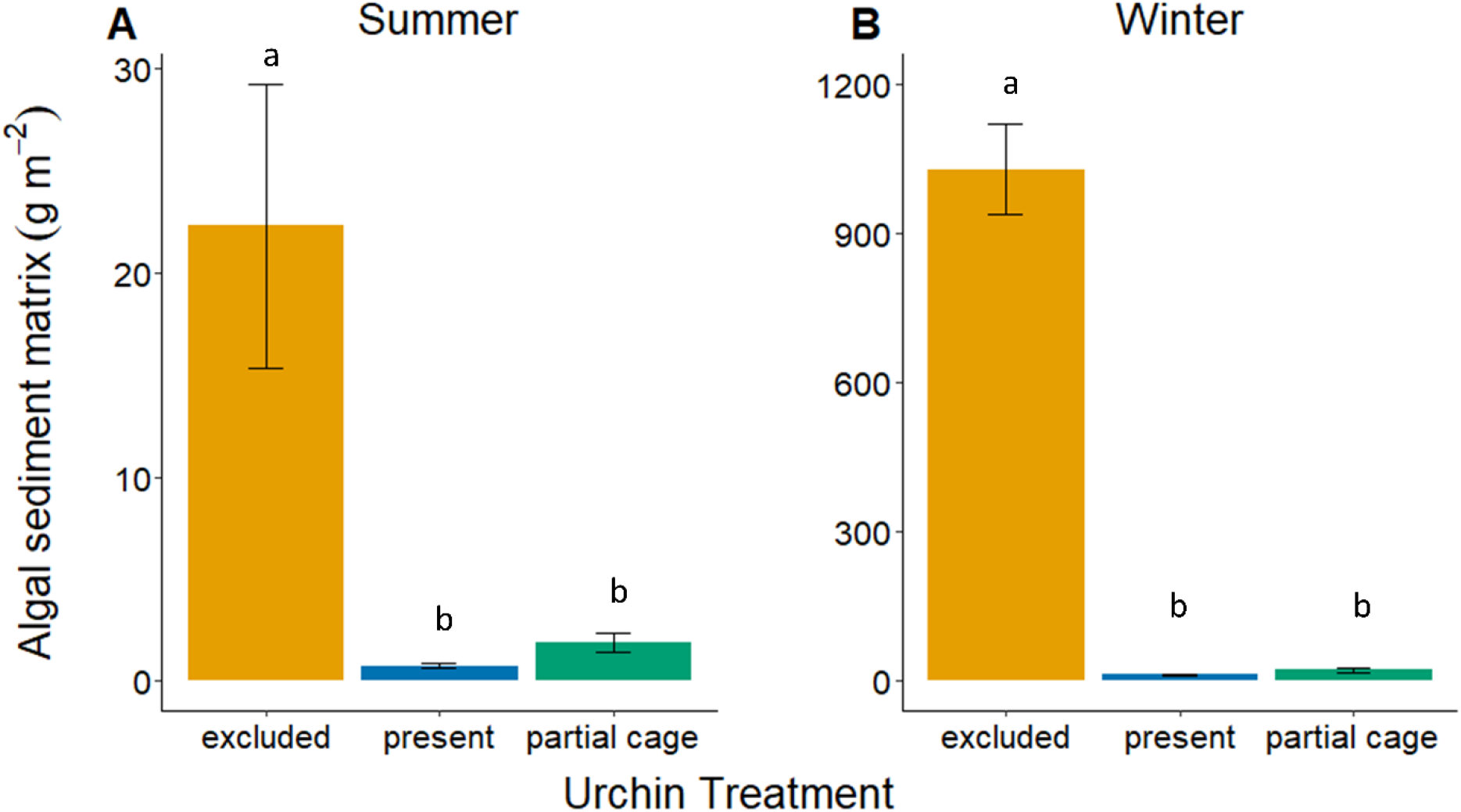
Growth of turf algae and accumulation of sediment (g × m^−2^ dry weight) when urchins are present or excluded for the (A) summer and (B) winter season. Values are the mean ± SE.

When urchins were present, *Porites sp.* had 100% survival, but this was reduced to 66.7% when urchins were excluded (Fig. S2). While overall survival of *P. decussata* was lower than *Porites sp.*, mortality was still higher when urchins were excluded than present (55.6% and 14.3%, respectively; Fig. S2). In contrast, *P. carnosus* suffered high mortality irrespective of urchin presence or absence. Consequently, new *P. carnosus* fragments were added to replace dead fragments on two separate occasions (Fig. S2).

Urchins increased the survival probability of *Porites sp.* coral fragments. When urchins were present, *Porites sp.* were 92% more likely to survive than when urchins were excluded (Cox test, Hazard Ratio (HR) = 0.08, 95% C.I. = 0.01 – 0.64, *p* = 0.018) and 84% more likely to survive in partial cages than when urchins were excluded (Cox test HR = 0.16, 95% C.I. = 0.03 – 0.80, *p* = 0.025). Consequently, the probability of *Porites sp.* surviving until the end of the experiment (393 days) was reduced from 90% to 30% when urchins were excluded (Fig. 3, Survival Analysis Log-Rank test, χ^2^ = 12.6 on 2 degrees of freedom, *p* < 0.002). By contrast, the presence of urchins had no effect on the survival probability (Cox test, *p* = 0.13, *p* = 0.60 for urchin presence and partial cages, respectively) or survival duration of *P. decussata* (Fig. 3, Log-Rank test, *p* = 0.260). Conversely, *P. carnosus* had a lower survival probability when urchins were present. When accounting for the death of fragments and the addition of new fragments, *P. carnosus* were 4.82 times more likely to die with urchins present, compared to when they were absent (Cox test, Hazard Ratio (HR) = 4.82, 95% C.I. = 1.49 – 15.6, *p* = 0.009). *P. carnosus* were also less likely to survive throughout the entire duration of the experiment when urchins were present (Fig. 3; Survival Analysis Log-Rank test, χ^2^ = 20.0 on 8 degrees of freedom, *p* < 0.01). However, when reducing the Survival Analysis model from 393 days (the duration of the entire experiment) to 163 days (the duration of the experiment after the last set of *P. carnosus* fragments were added) the probably of *P. carnosus* surviving until the end of the experiment (the remaining 163 days) increases to 71.4% ± 12.1% SE when urchins were present.

**Figure 3.**
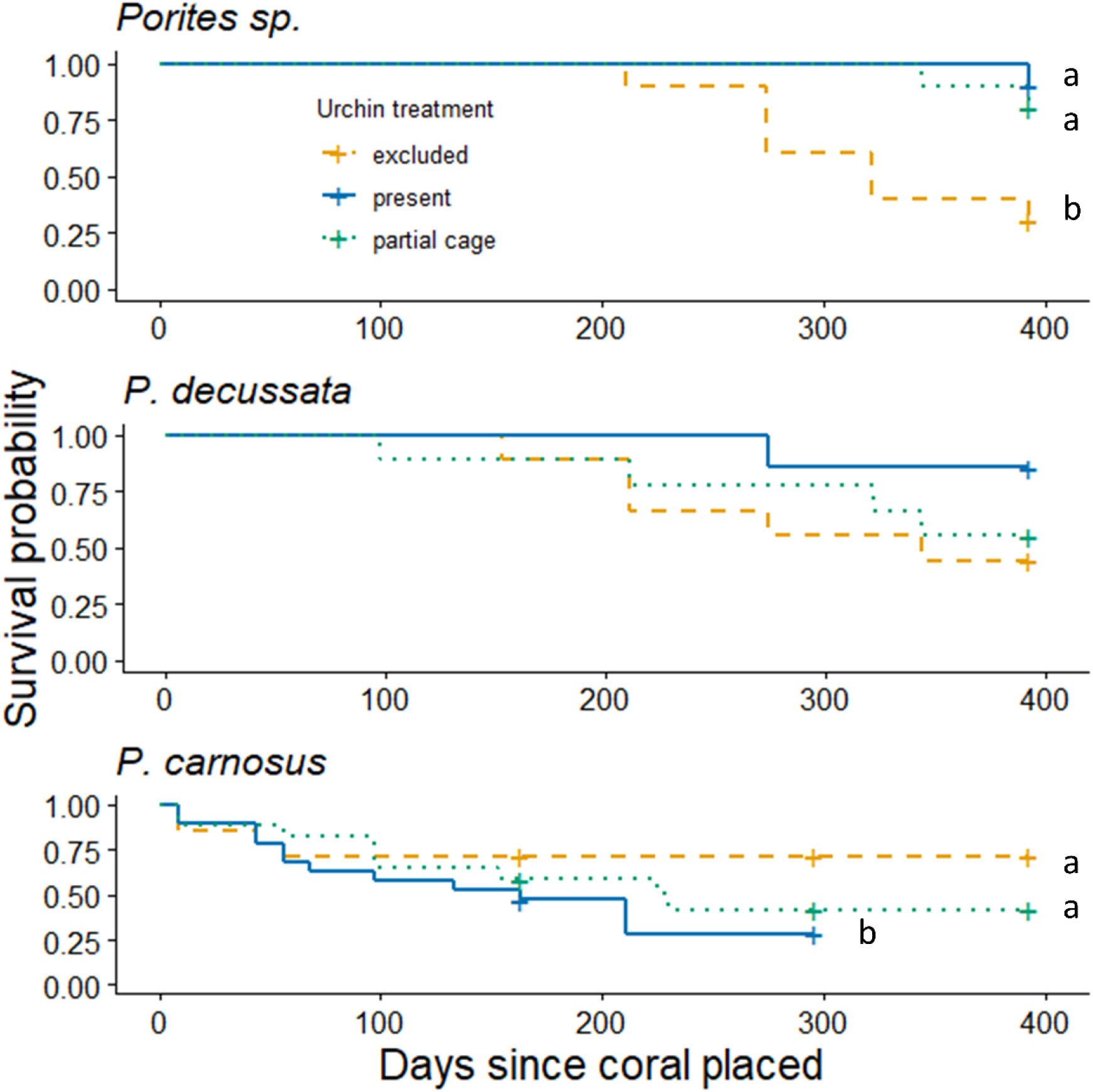
The probability of each species of coral surviving until the end of the experiment when urchins are absent (yellow), present, (blue) absent (green), and under procedural control (blue). Kaplan-Meier plot, crosses indicated censored data points, coral fragments that did not die before the conclusion of the experiment. Letters represent significant differences amongst treatments.

*Porites sp.* and *P. decussata* had similar growth rates regardless of urchin presence or absence: in both cases they grew throughout the year (Fig. 4). There was no difference in buoyant weight of surviving fragments of both *Porites sp.* and *P. decussata* when urchins were excluded relative to when urchins were present (One-way ANOVA, *F* _2, 16_ = 0.58, *p* = 0.571 and *F* _2, 12_ = 1.094, *p* = 0.366, respectively). Though *Porites sp.* grew at similar rates when urchins were excluded, they were more likely to survive, and therefore have greater net growth when urchins are present. *P. decussata* had similar, though not significant survival and growth trends as *Porites sp.* and may contribute to overall net growth. In contrast, when urchins were present *P. carnosus* decreased in weight due to bioerosion (Fig 4.; *F* _2, 21_ = 10.4, *p* < 0.001).

**Figure 4.**
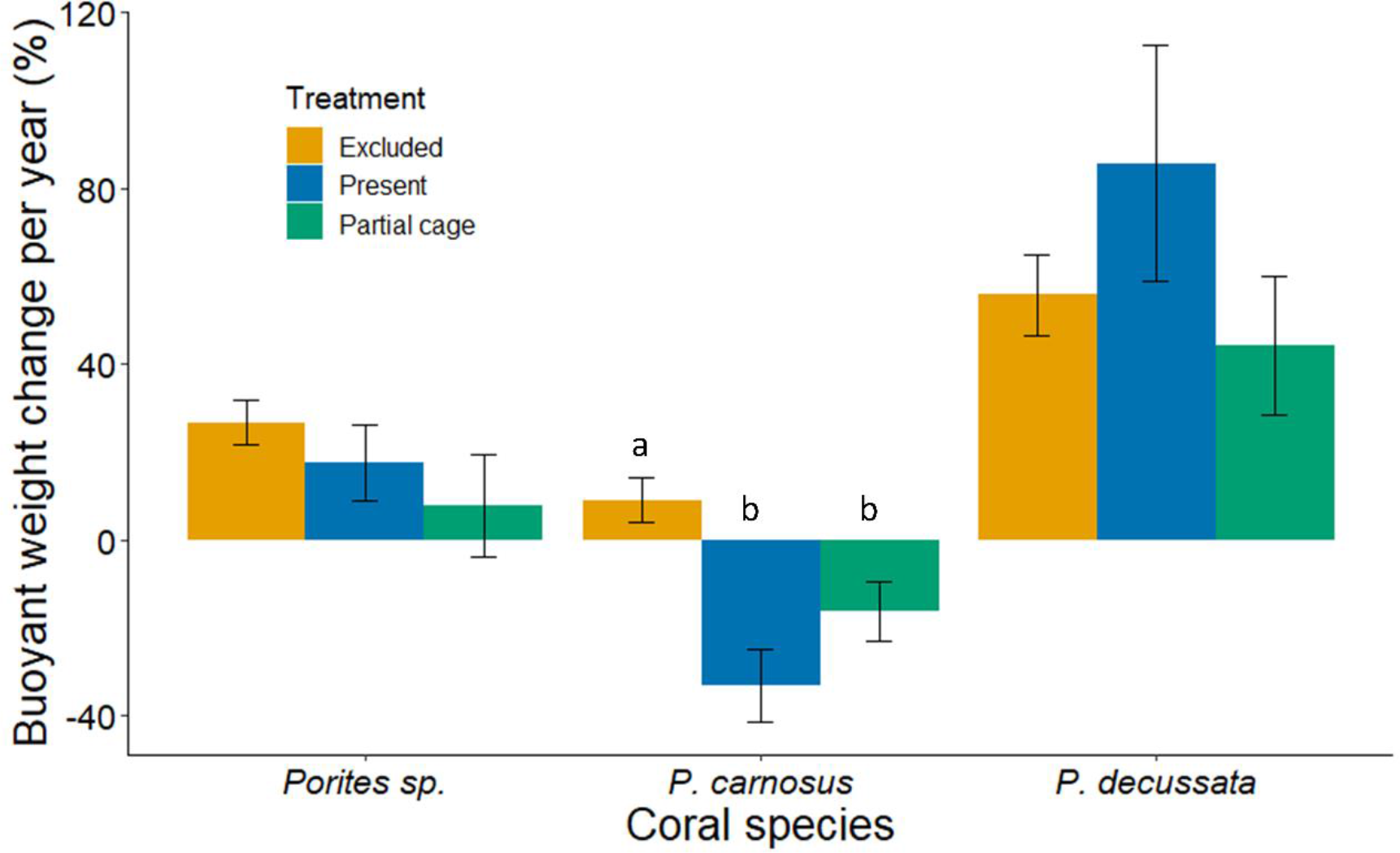
Percent change in the buoyant weight of corals per year when urchins were excluded, present, and present in partially caged plots. Only corals that were still alive at the end of the experiment were used for these analyses. Values are mean percent ± SE. Asterisks represent significant differences amongst treatments.

The presence of urchins increased bioerosion of coral skeleton cores by 4.5 times in the summer and 3.1 times in the winter (Fig. 5; *F* _2, 23_ = 59.83, *p* < 0.001, *F* _2, 24_ = 9.13, *p* < 0.002, respectively). There were no statistical differences between the partially caged and urchin present plots during the summer and winter (Tukey HSD *p* = 0.46, *p* = 0.059, respectively). When urchins were excluded, the coral skeleton cores accumulated turf, sediment and macroalgae like the settlement tiles.

**Figure 5.**
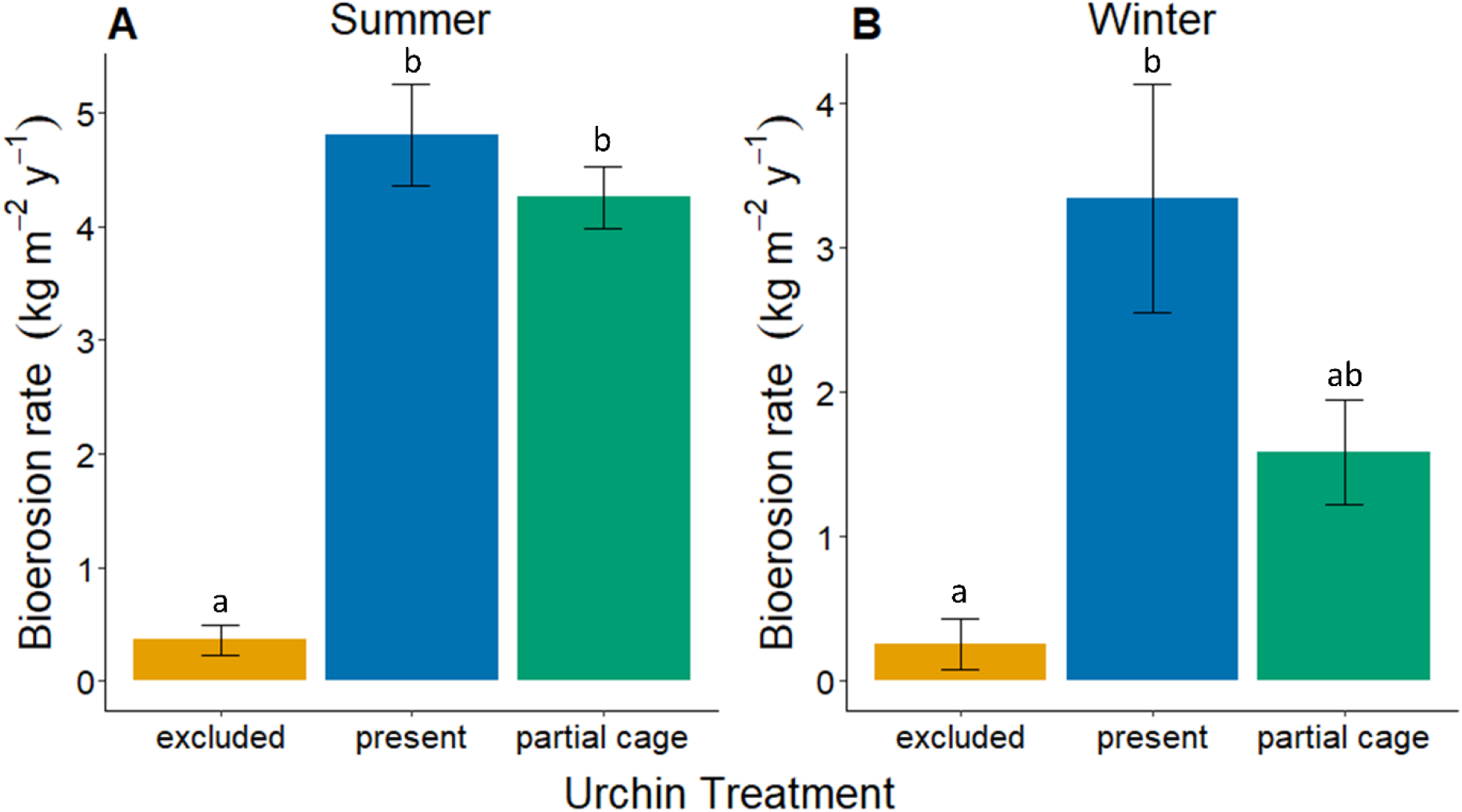
The rates of bioerosion of coral skeleton core disks by the urchin *Diadema setosum* for (A) summer and (B) winter. Values are mean ± SE. Letters represent significant differences between urchin treatments.

## Discussion

Urchins are important contributors to the health and function of coral reefs. They can prevent overgrowth of hard corals by macroalgae and maintain space bare of algae which increases coral larvae settlement (Carpenter and Edmunds 2006, Hughes et al. 2007). Yet, eutrophic conditions are generally thought to cause a shift from coral to macroalgae dominated reefs as algal growth overwhelms herbivory. Here, we show that even in a heavily developed and eutrophic system, urchins reduce algal-sediment matrix, increase CCA coverage, and subsequently can increase coral survival. When urchins were present, they reduced the benthic coverage of algae and trapped sediment by up to 14-fold, corresponding to an 80-fold decrease in algal biomass and sediment mass. This reduction in algae and sediment accumulation is crucial for coral survival as algae can overgrow coral and prevent recruitment (Samarrco 1980, Edmunds and Carpenter 2001, Carpenter and Edmunds 2006, Hughes et al. 2007, Nozawa et al. 2020), and sediment can smother coral, both causing coral death (Weber et al. 2006, Risk and Edinger 2011, Weber et al. 2012). Therefore, if herbivores, and in this case the long-spined sea urchin, *D. setosum*, are absent from large areas of coral based ecosystems for a sustained period, a phase shift from coral to algae generally occurs (Carpenter and Edmunds 2006, Hughes et al. 2007, Reverter et al. 2020). While there was a substantial overall positive effect of urchins in the system, the effects on individual corals were more nuanced and species specific than generally reported. Given that urchins reduced algal growth and sediment accumulation, we expected coral growth to be enhanced by the presence of urchins. Instead, we saw no difference in growth for two of the three species, *P. decussata* and *Porites sp.*, and negative growth, or bioerosion, for *P. carnosus*. Such nuanced outcomes are not uncommon. For example, several coral species increased in area by 20% when two herbivorous fishes were present, yet when only one fish was present or both were excluded, corals decreased in area (were bioeroded) by up to 30% (Burkepile and Hay 2008). Such species-specific responses are also commonly observed in corals with natural herbivore communities: for example, *Porites astreoides* grew four times faster when urchins and fishes were present, yet *Siderastrea siderea* was unaffected by herbivore exclusion (Lirman 2001). Therefore, not all corals respond similarly to herbivores and not all herbivores have the same affects on corals growth. Importantly, however, survival rates of corals are essential to incorporate in assessments, as the presence of herbivores tends to increase survival for most species (this study; Lirman 2001, Burkepile and Hay 2008, Knoester et al. 2019). Therefore, while herbivores may cause little difference in coral growth, they increase coral survival, which leads to net growth of coral reefs.

Urchins have been associated with increased health of coral reefs and the survival of coral colonies by reducing macroalgae and limiting coral overgrowth (Edmunds and Carpenter 2001, Carpenter and Edmunds 2006, Hughes et al. 2007). We found that the effect of urchins on coral survival is species-dependant; survival of *Porites sp.* increased, *P. carnosus* decreased, and there was no effect on *P. decussata.* Though *Porites sp.* and *P. decussata* settled well after transplantation, *P. carnosus* seemed to immediately suffer mortality upon transplant which could be because this species is more susceptible to mortality under stressful conditions (e.g. eutrophication; Fabricius 2005, Cybulski et al. 2020). Survival in other coral species have been shown to increase when systems are healthier and have functionally redundant herbivores. For example, *Orbicella faveolata.* have a two-fold increase in survival (Lirman 2001) and *Acropora verweyi* a 10% increase in survival (Knoester et al. 2019) when compared to those when herbivores were excluded. Similarly, in an inclusion experiment in the Florida Keys, coral colonies had 100% survival when two herbivore species were present, but only 75% survival when both were excluded (Burkepile and Hay 2008) further emphasizing the importance of having a multiple herbivore species in an ecosystem. This is especially important in eutrophic systems where algae grow at increased rates and the compensatory feeding of herbivores is less able to reduce macroalgal dominance (Littler et al. 2006, Ghedini et al. 2015).

Though coral growth rates were lower than we expected when urchins were present, the increase in CCA cover is an encouraging sign for the long-term health of coral communities as CCA promotes coral recruitment, increases structural integrity of reefs, and deters macroalgal settlement. CCA cover is well documented as being important as a chemical cue for coral larval settlement and recruitment (Heyward and Negri 1999, Harrington et al. 2004, Goméz-Lemos et al. 2018). In fact, several studies have found that CCA increased coral settlement and metamorphosis from 20% to 85% over other substrata (Heyward and Negri 1999, Goméz-Lemos et al. 2018). As we found urchins increased CCA cover by 60% in both summer and winter, we would expect to see increased coral settlement rates over longer periods, though we did not test for this. Further, CCA also acts a binding agent, or cement, helping reinforce the strength and integrity of coral reefs (Littler and Littler 2013, Weiss and Martindale 2017). As CCA grows over dead coral skeletons and between corals, it reinforces the skeletons and increases resistance to physical disturbances (Littler and Littler 2013). Over time, the aragonite in thick CCA crust dissolves and re-precipitates as dolomite, a substantially harder mineral than the aragonite in coral skeleton (Nash et al. 2011, Nash et al. 2013) and is associated with stronger reef framework (Weiss and Martindale, 2017). Further, urchin grazing activity results in grazer-resistant CCA (Davis 2009), which could reduce bioerosion and further increase coral structural integrity. Lastly, CCA has be found to deter the settlement and growth of macroalgae (Vermeij et al. 2011, Gómez-Lemos and Diaz-Pulido 2017). For example, *Padina boergesenii* spore settlement was nine times lower on live CCA versus dead CCA (Gómez-Lemos and Diaz-Pulido 2017), whereas *Ulva fasciata* growth was supressed by 55% when grown with CCA (Vermeij et al. 2011). As we found that urchin presence greatly increases CCA coverage, the secondary benefits of CCA may contribute to healthier coral communities in Hong Kong.

The accumulation of sediment can negatively affect corals in several ways: direct smothering of corals; decreasing light availability which limits photosynthesis; an increased metabolic demand on corals to shed the sediment; and microbial processes such as respiration, fermentation and desulfurylation (Risk and Edinger 2011, Weber et al. 2012). As sediment either accumulates on corals or is suspended in the water column, the amount of light that reaches corals is reduced, decreasing the photosynthetic yield of corals and can lead to photosynthetic stress and bleaching by reducing net productivity (Weber et al. 2006, Risk and Edinger 2011). Furthermore, when sediment accumulates on corals, the colonies actively remove sediment through tentacle waving, cilia action or mucus production, all of which increase coral metabolic demand while reducing feeding and photosynthetic capabilities (Risk and Edinger 2011). Lastly, sediment-derived microbial respiration creates anoxic conditions and can kill coral tissues in as little as 15 hours (Weber et al. 2006, Weber et al. 2012). In systems which have become eutrophic, where microbial communities have shifted to more pathogenic and sulphate-reducing organisms such as Hong Kong (Chen et al. 2019), it is possible that this microbial degradation could be exacerbated. Decreased photosynthetic capability, increased metabolic costs, and anoxic conditions caused by sediment buildup all have negative impacts on corals and can decrease their potential survival capabilities. As we have shown here, urchins can greatly reduce algal sediment matrices, and are an important part of coral resilience.

While urchins can scrape away algae which increases CCA cover and provides settlement areas for juvenile corals, overgrazing can lead to bioerosion of the coral skeleton (Glynn 2015, Alvarado et al. 2017) and can be detrimental to a coral ecosystem when the erosion rate is greater than accretion (Alvarado et al. 2016). Bioerosion generally increases only after conditions lead to coral tissue death, leading to algal growth on the skeleton and favouring bioeroders over corals (Barkley et al. 2015, Glynn, 2015). In a healthy reef or coral community replete with an intact trophic web, bioerosion by urchins is minimal (Bellwood et al. 2004, Alvarado et al. 2017). When ecosystems are degraded, however, a system imbalance can drive increased bioerosion rates, but how this manifests depends on the local urchin species, population density, and size of the individuals (Glynn 2015, Alvarado et al. 2017). Urchin bioerosion rates can range from 0.07 – 10.4 kg · m^−2^ · yr^−1^, with urchin densities varying from less than 1 to more than 100 individuals m^−2^ (Glynn 2015). We found that when urchins were present, bioerosion ranged between 3.3 – 4.8 kg · m^−2^ · yr^−1^ for winter and summer, respectively. m^−2^ · yr^−1^ meaning that net bioerosion by urchins was between 3.0 – 4.4 kg · m^−2^ · yr^−1^ in this system. These rates are moderate when compared to other reef systems globally. Importantly, urchin bioerosion is often measured by extracting fecal material, separating out all calcium carbonate, and determining gut turn-over rates to estimate bioerosion (Dumont et al. 2013, Alvarado et al. 2016). In contrast, we calculated bioerosion from the loss of skeletal CaCO_3_ when urchins were present and absent, providing a direct (and more accurate) measure.

Herbivory is widely accepted as being beneficial to overall coral reef health, though the reality is much more nuanced, species-specific, and dependent on the overall state of the system. As we have shown, herbivory has positive effects on coral communities through increased CCA cover, which promotes coral settlement, strengthens reefs, and deters macroalgal settlement, and overall increased coral survival. Several studies have also demonstrated faster coral growth when herbivores are present, though ours did not. Perhaps most importantly, herbivory substantially decreases algal cover and sediment accumulation, both of which kill corals and suppresses growth and recruitment if left unchecked. Yet not all corals react the same way to herbivory. *P. carnosus*, for instance, seem to be more susceptible to bioerosion when either alive or dead (Dumont et al. 2013, Qiu et al. 2014, this study). We therefore caution that herbivory has the potential to be deleterious to some corals when overfishing has led to grazer densities that are unnaturally high, but at natural densities the benefits of herbivores far outweigh any potential negatives. Importantly, degraded or highly disturbed ecosystems (i.e., overfished, eutrophic, increased terrestrial sediment run-off, bleached, etc.) can be strengthened or even restored through the positive effects of herbivores. Ultimately, it is necessary to take all aspects of herbivory into account, including coral and herbivore community diversity and composition, when evaluating the role of herbivory in coral reef ecosystems.

## Acknowledgements

We would like to thank A. Anand, R. Cheung, R. Gotama, A. Hemraj, T. Kim, J. Minuti, and V. Yu for their support in the construction, deployment, surveys and clean up of the experiment. This study research was supported by an Agriculture, Fisheries and Conservation Department (AFCD) Hong Kong contract (AFCD/SQ/3/16/C), the Environment and Conservation Fund #67/2016 and a Collaborative Research Fund award (#C7013-19G) from the Hong Kong Research Grants Council to BDR and DMB. We would also like to thank the Director of the Agriculture, Fisheries and Conservation Department for permission to publish this paper. This manuscript is contribution # XXXX to MarineGEO.

